# Detection of SARS-CoV-2 Omicron variant (B.1.1.529) infection of white-tailed deer

**DOI:** 10.1101/2022.02.04.479189

**Authors:** Kurt J. Vandegrift, Michele Yon, Meera Surendran-Nair, Abhinay Gontu, Saranya Amirthalingam, Ruth H. Nissly, Nicole Levine, Tod Stuber, Anthony J. DeNicola, Jason R. Boulanger, Nathan Kotschwar, Sarah Grimké Aucoin, Richard Simon, Katrina Toal, Randall J. Olsen, James J. Davis, Dashzeveg Bold, Natasha N. Gaudreault, Juergen A. Richt, James M. Musser, Peter J. Hudson, Vivek Kapur, Suresh V. Kuchipudi

## Abstract

White-tailed deer (*Odocoileus virginianus*) are highly susceptible to infection by SARS-CoV-2, with multiple reports of widespread spillover of virus from humans to free-living deer. While the recently emerged SARS-CoV-2 B.1.1.529 Omicron variant of concern (VoC) has been shown to be notably more transmissible amongst humans, its ability to cause infection and spillover to non-human animals remains a challenge of concern. We found that 19 of the 131 (14.5%; 95% CI: 0.10–0.22) white-tailed deer opportunistically sampled on Staten Island, New York, between December 12, 2021, and January 31, 2022, were positive for SARS-CoV-2 specific serum antibodies using a surrogate virus neutralization assay, indicating prior exposure. The results also revealed strong evidence of age-dependence in antibody prevalence. A significantly (χ^2^, p < 0.001) greater proportion of yearling deer possessed neutralizing antibodies as compared with fawns (OR=12.7; 95% CI 4–37.5). Importantly, SARS-CoV-2 nucleic acid was detected in nasal swabs from seven of 68 (10.29%; 95% CI: 0.0–0.20) of the sampled deer, and whole-genome sequencing identified the SARS-CoV-2 Omicron VoC (B.1.1.529) is circulating amongst the white-tailed deer on Staten Island. Phylogenetic analyses revealed the deer Omicron sequences clustered closely with other, recently reported Omicron sequences recovered from infected humans in New York City and elsewhere, consistent with human to deer spillover. Interestingly, one individual deer was positive for viral RNA and had a high level of neutralizing antibodies, suggesting either rapid serological conversion during an ongoing infection or a “breakthrough” infection in a previously exposed animal. Together, our findings show that the SARS-CoV-2 B.1.1.529 Omicron VoC can infect white-tailed deer and highlights an urgent need for comprehensive surveillance of susceptible animal species to identify ecological transmission networks and better assess the potential risks of spillback to humans.

**Key Findings:** These studies provide strong evidence of infection of free-living white-tailed deer with the SARS-CoV-2 B.1.1.529 Omicron variant of concern on Staten Island, New York, and highlight an urgent need for investigations on human-to-animal-to-human spillovers/spillbacks as well as on better defining the expanding host-range of SARS-CoV-2 in non-human animals and the environment.

## Main Text

Recent investigations have established that severe acute respiratory syndrome coronavirus-2 (SARS-CoV-2) infects a wide range of non-human animal hosts^1-3^. SARS□CoV□2 infections have been documented in several animal hosts, including farmed mink, companion animals (e.g., cats, dogs, ferrets), and zoo animals (e.g., tigers, lions, cougars, snow leopards, gorillas, otters, and hippopotamus)^4,5^. White-tailed deer (*Odocoileus virginianus*) are highly susceptible to SARS-CoV-2 as evident by the widespread natural infections in Iowa^6^ and Ohio^7^, together with experimental infection studies with ancestral SARS-CoV-2 or the Alpha variant of SARS-CoV-2^8,9^. More recently, SARS-CoV-2 infection of deer from multiple states in the USA has been confirmed^10^, further heightening concerns for the potential of deer to serve as a reservoir of SARS-CoV-2. Recent SARS-CoV-2 variants of concern (VoCs) such as Delta and Omicron appear to be more highly transmissible between humans than previously described strains and variants^11,12^. While there are reports of zooanthroponotic spillover of the SARS-CoV-2 Delta VoC into multiple species including cats, dogs, pumas and lions in a zoo in South Africa^13^, free-living deer across many US states and Syrian hamsters in pet shops in Hong Kong^14^, the spillover of the Omicron VoC to non-human animal species has not yet been documented. Given the lack of clarity on the origins of Omicron, and whether it emerged from a chronically infected human host, silently spread in a cryptic human population, or emerged from a yet unknown non-human animal population (e.g., rodents^15^), a current research priority is to understand the host range and evolutionary trajectories of Omicron in potential animal reservoirs.

Here we report spillover of SARS-CoV-2 Omicron VoC infections into the white-tailed deer population inhabiting Staten Island, New York. To our knowledge, this is the first report of Omicron infection in a wildlife species. The identification of a white-tailed deer with evidence of shedding viral RNA in nasal secretions in the presence of circulating serum neutralizing antibodies raises the intriguing possibility that deer, as with humans, may also experience breakthrough infections and reinfections with SARS-CoV-2. Taken together, our studies highlight the urgent need for better assessing the risks associated with the expanding host-range and understanding the evolutionary trajectories of Omicron and other SARS-CoV-2 strains and variants in non-human animal reservoirs and the environment.

## Methods

### Samples

Deer were opportunistically darted and anesthetized by wildlife biologists during an ongoing deer sterilization study implemented by the City of New York Parks & Recreation (NYC Parks). Once darted, animals were tracked and processed at a sampling site. Anesthetized animals were given ear tags, sampled, and released. The GPS location of the sampling site for each individual was recorded, and the animal’s sex and age were determined. Blood samples were collected from the jugular vein into serum separator tubes and the serum was frozen at - 20°C until testing. Nasal and palatine tonsil swabs were obtained using Copan floQ swabs (Copan Diagnostics Inc.) Swabs were placed directly in Universal Transport Media (UTM). Samples were submitted to the Penn State Animal Diagnostic Laboratory (ADL) for diagnostic testing and were subsequently used in this study.

### Surrogate Virus Neutralization Test (sVNT)

Serum samples were screened using a surrogate virus neutralization test (sVNT) assay that was previously validated for detecting SARS-CoV-2 antibodies in deer^16^. The sVNT assay uses cPass™ technology (Genscript) and detects total neutralizing antibodies measured as percent inhibition^17^. Animals with a percent inhibition above 30% were considered positive.

### Indirect ELISA assays for RBD and N protein

In-house developed indirect ELISA (iELISA) assays were used to determine antibody titers against the SARS-CoV-2 Spike (S) receptor-binding domain (RBD) and nucleocapsid (N) protein. Briefly, nucleotide sequences encoding the nucleocapsid N protein and the RBD of the S protein of the SARS-CoV-2 isolate Wuhan-Hu-1 (GenBank accession #: MT380725.1) were used for constructing plasmids for protein expression. The N and RBD genes, plus two streptavidin tags, were cloned into the mammalian expression vector pHL, and the recombinant proteins were produced in HEK (human embryo kidney) 293 cells as described previously^9^. Using a panel of known positive and pre-pandemic negative deer sera, cutoffs for the iELISA were established.

### RNA Extraction and RT-PCR for SARS-CoV-2 detection

Processing of swab samples and real-time RT-PCR were conducted in a Clinical Laboratory Improvement Amendment (CLIA)-approved laboratory. RNA was extracted from 400 µL of swab samples using a KingFisher Flex machine (ThermoFisher Scientific) with the MagMAX Viral/Pathogen extraction kit (ThermoFisher Scientific) following the manufacturer’s instructions. The presence of SARS-CoV-2 viral RNA was tested using the OPTI Medical SARS-CoV-2 RT-PCR kit, which is a highly sensitive assay targeting nucleocapsid (N) gene with a limit of detection of 0.36 copies/µL^18,19^. RT-PCR assays were carried out on an ABI 7500 Fast instrument (ThermoFisher Scientific). The internal control RNase P (RP) was utilized to confirm samples were not contaminated with human tissue or fluids during harvesting or processing. All samples were found to be negative for the presence of the human RP gene by RT-PCR. Samples were also tested using a TaqPath kit (ThermoFisher Scientific) that targets the SARS-CoV-2 ORF1ab, N gene, and S gene^19,20^ as the first screen for Omicron, which typically presents as an S gene drop out in these assays^21^.

### SARS-CoV-2 Genome Sequencing

Total RNA extracted from swab samples was used for whole genome sequencing of SARS-CoV-2 as previously described^6,22-25^ and the sequencing libraries were prepared according to version 4.1 of the ARTIC nCoV-2019 protocol^26^. We used a semi-automated workflow that employed BioMek i7 liquid-handling workstations (Beckman Coulter Life Sciences) and MANTIS automated liquid handlers (FORMULATRIX). Using a NovaSeq 6000 instrument (Illumina) we generated short sequence reads to ensure a very high depth of coverage. Sequencing libraries were prepared in duplicate and sequenced with an SP 300 cycle reagent kit.

### SARS-CoV-2 Genome Sequence Analysis and Identification of Variants

Viral genomes were assembled using the BV-BRC SARS-CoV2 assembly service^27,28^, which uses a pipeline that is similar to One Codex SARS-CoV-2 variant-calling pipeline^29^. Briefly, the pipeline uses seqtk version 1.3-r116 for sequence trimming^30^, minimap version 2.1^31^ for aligning reads against reference genome Wuhan-Hu-1 NC_045512.2^32,33^, samtools version 1.11^34^ for sequence and file manipulation^35^, and iVar version 1.2.2^36^ for primer trimming and variant calling^37^. To increase stringency, the minimum read depth for assemblies (based on samtools mpileup) was set at three to determine consensus. Genetic lineages, variants being monitored, and VoCs were identified and designated by Pangolin version 3.1.11 (14)^22^ with pangoLEARN module 2021-08-024. Single Nucleotide Polymorphisms (SNPs) were identified using the vSNP (18) SNP analysis program.

### Data Analysis and Visualization

Serology data was analyzed using the z-test for differences in proportions and visualized using GraphPad Prism version 9.0 for macOS (GraphPad Software, San Diego, California USA). QGIS (geographic information system) mapping software version 3.16.10 was used to visually portray the geographic location of the white-tailed deer that were sampled ^38^. For age-group comparisons, deer with estimated ages <12 months were categorized as fawns; deer between 12-29 months were categorized as “yearlings”; and deer ≥30 months were categorized as adults.

## Results and Discussion

### Serological evidence for SARS-CoV-2 exposure of white-tailed deer on Staten Island, New York

To determine if white-tailed deer on Staten Island have previously been exposed to SARS-CoV-2, serum samples from 131 individual deer collected between December 12, 2021 and January 31, 2022 as part of an ongoing population control study and disease surveillance program were examined for the presence of anti-SARS-CoV-2 neutralizing antibodies using sVNT^16^. As a consequence of the sampling design targeting males, the vast majority of the serum samples were from males (*n* = 116; 88.5%) with an age distribution heavily skewed toward the younger age classes. Eighty-two fawns constituted 62.6% of the sample and 33 yearling deer made up 25.2%, while only 16 of the 131 individuals (12.2%) were considered adults. The results show that 19 of 131 (14.5%; 95% CI: 0.1–0.22) serum samples were positive for SARS-CoV-2 exposure with the surrogate virus neutralization test. Viral inhibition in the positive samples ranged from 33.2% to 97%, with a median value of 70.9% (Table 1, Figure 1).

**Figure 1.**
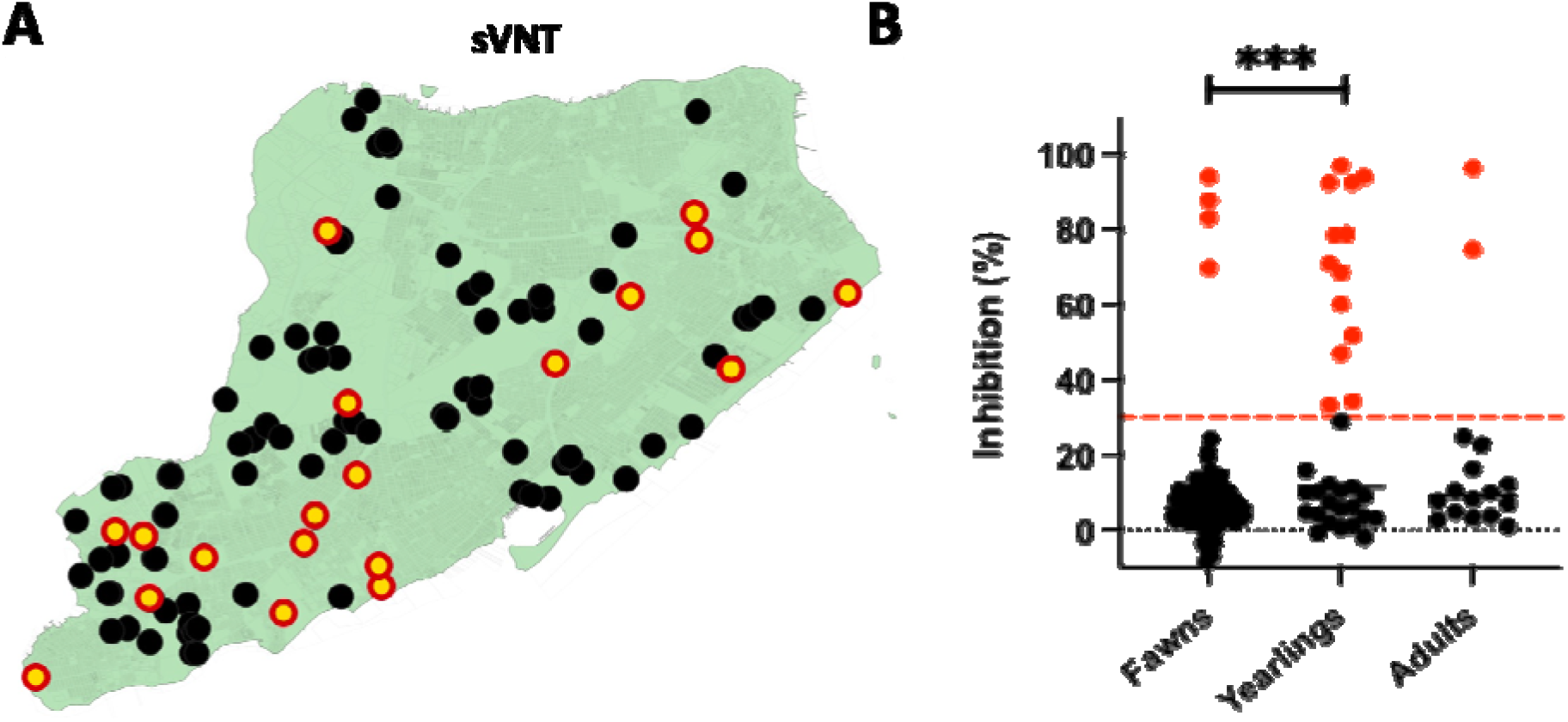
SARS-CoV-2 serological reactivity status of white-tailed deer on Staten Island, New York between December 12, 2021 and January 30, 2022. (A) Spatial distribution of sites of collection of serum samples from white-tailed deer for assessment of serological reactivity to and neutralization of SARS-CoV-2. Red circles with yellow centers indicate positive detection and black filled circles indicate seronegative status. (B) Age-group stratified serum virus neutralization activity in white-tailed deer on Staten Island, NY. Serum neutralization was determined using the commercially available SARS-CoV-2 Serum Virus Neutralization Assay (sVNT; Genscript cPass) and age-group percent inhibition determined. \*\*\**p*<0.0001.

The proportion of positive animals is not different than the findings of Chandler *et al*.^16^, who identified nine of 29 (31%; 95% CI 0.17, 0.49) white-tailed deer from two New York counties (Suffolk and Onondaga) seropositive to SARS-CoV-2 in 2021. The results also reveal age-dependent variation in the distribution of neutralizing antibodies in the sampled white-tailed deer, with a significantly greater (z = 4.2; p < 0.01) proportion of antibody positive individuals represented by yearlings (39.4%; 95% CI: 0.25–0.56) as compared with fawns (4.9%; 95% CI: 0.02–0.11) (Figure 1). Only two of the 16 (12.5%; 95% CI: 0.02–0.40) adults we sampled had evidence of antibodies and thus the small number of samples in this age group precludes robust comparisons.

Chandler *et al*^*16*^ showed 82% of antibody seroprevalence in yearling (∼1.5 year-old) deer, which was the highest among all the sampled age groups^16^, and Palermo and colleagues^39^ also noted a similarly elevated seroprevalence in yearlings as compared with other age groups in Texas. While the exact reasons for the observed higher SARS-CoV-2 specific antibody prevalence in yearlings is unknown, the behavioral characteristics and social stressors on male deer transitioning from their maternal home range might play a role. Regardless, further studies on the impacts of social networks as well as life stage and seasonal behaviors (such as the rut) on the susceptibility and transmission dynamics of SARS-CoV-2 in white-tailed deer are needed to fill important knowledge gaps. Importantly, while our results suggest an intriguing age-dependent variation in observed seroprevalence in white-tailed deer, SARS-CoV-2 is likely in an epizootic and not an enzootic stage in white-tailed deer, and hence estimations of infection rates or the basic reproduction number, *R*_*0*_, are not feasible from the current estimates and require further, preferably longitudinal, investigations. Future investigations are also needed to assess whether the observed higher seroconversion in yearlings as compared with fawns or adults may be of use in the rationale design of targeted surveillance approaches.

### Molecular and genetic identification of SARS-CoV-2 Omicron VoC in nasal and tonsillar swabs from white-tailed deer in Staten Island, New York

There is prior antibody^16^ and RT-PCR^10^ evidence of SARS-CoV-2 exposure of white-tailed deer in New York. However, it is not clear whether deer previously exposed to SARS-CoV-2 might be reinfected and whether there is continued SARS-CoV-2 spillover infection of deer in urban settings after the emergence of the Omicron VoC. To address this, we used a well-validated RT-PCR assay to detect the presence of

All retested samples showed amplification of orf1a/b and S gene dropout in the TaqPath multiplex assay, suggestive of Omicron VoC as previously described^21^. However, we also observed an unusual N-gene dropout or Ct value shift in most of the samples with the TaqPath assay.

To determine the identity and genetic relatedness of the circulating strains, whole-genome sequencing (WGS) was applied on four of the seven positive samples (tags 102–104 and 2067) using a recently described pipeline^6^. Sequencing of 3 RT-PCR positive samples (numbers 2089, 2108 and 2125) is currently underway. The analysis confirmed that all four positive samples were represented by the Omicron lineage that was widespread and the dominant circulating lineage (nearly 90%) amongst humans in New York City starting during the latter part of December of 2021 through January 2022^40^. To our knowledge, this is the first report of the Omicron VoC of SARS-CoV-2 from white-tailed deer or any other wildlife. This finding indicates that, together with reports of experimental and natural infections of white-tailed deer with ancestral SARS-CoV-2 or Alpha VoC, as well as natural infection with the Delta and Gamma VoCs, deer are also susceptible to the Omicron VoC.

Phylogenetic analyses of the whole genome sequences of these newly identified Omicron sequences from white-tailed deer were performed with the vSNP pipeline followed by visualization using iToL^41^ (Figure 2B). The analysis shows that these sequences cluster closely with recently reported Omicron sequences from humans in New York City as well as with those reported from environmental sources in Austria but are quite distinct from the previously described isolates recovered form free-living deer in Iowa, Ohio^6,7^ and from 13 other US states that have recently been deposited in GISAID^42^ (Accessions EPI_ISL_9347708 through EPI_ISL_9347728 and EPI_ISL_9388140 through EPI_ISL_9388144) (Figure 2B and Supplemental Table 2). The closely clustered viral genomes from samples collected from deer 102, 103 and 104 that were all sampled on the same date (January 6, 2022) and the same location had between 5–6 SNPs consistent with a common point source of infection or recent transmission between individual animals. The viral RNA recovered from the fawn 2067 which was sampled a day later from a different location revealed 9–12 SNPs compared with those from fawns 102, 103 and 104, suggesting a potentially different point source of infection. However, confidence in the robustness of inferences that can be drawn from these data regarding point sources of infection or transmission is limited because the currently available dataset is very small, and it is likely missing a large number of sequences. We also note that because of the overall comparatively low evolutionary rate relative to generational interval, and similar length of window for transmission of SARS-CoV-2 relative to the incubation period in deer, the reliable reconstruction of statistically robust transmission networks will require much more representative and frequent sampling.

**Figure 2.**
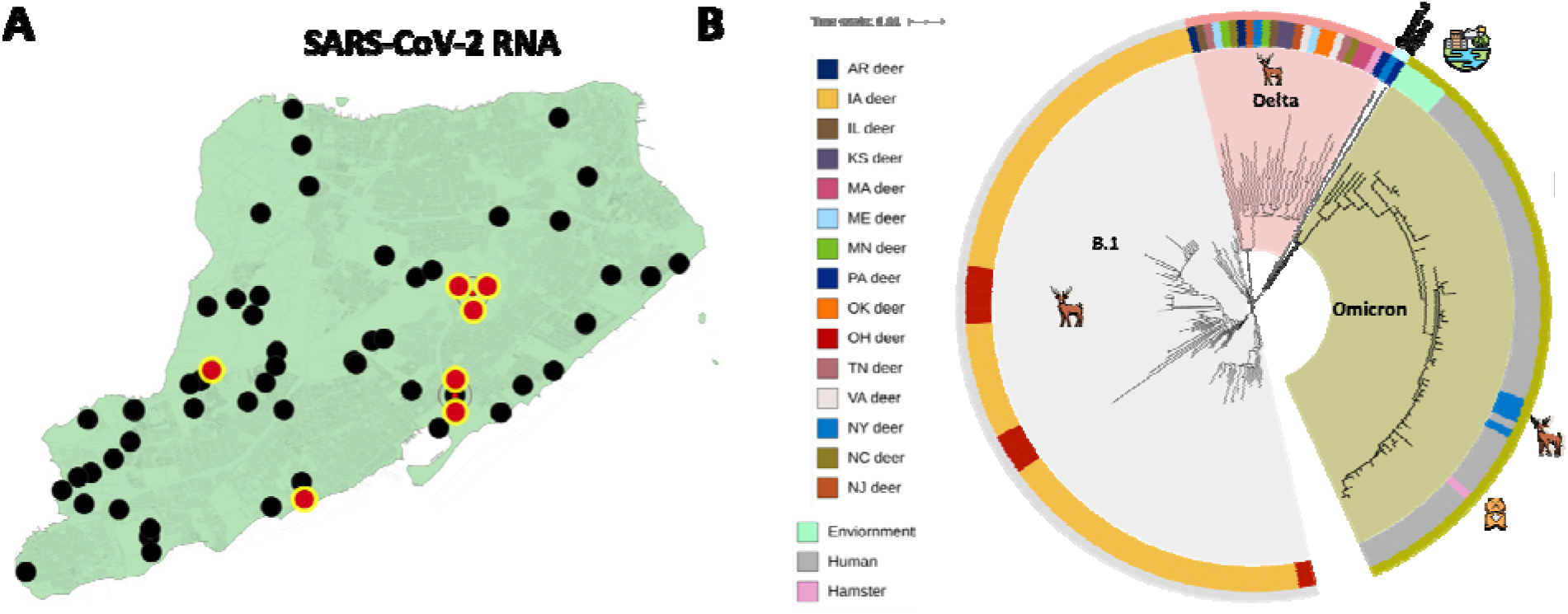
Distribution and whole-genome single nucleotide polymorphism (SNP)-based phytogenies of SARS-CoV-2 recovered from white-tailed deer on Staten Island. New York. (A) The spatial distribution of the collection sites of nasal swabs from white-tailed deer which were tested for the presence of SARS-CoV-2 viral RNA. Magenta circles with blue highlights show sites where swabs positive for SARS-CoV-2 viral RNA were sampled and black filled circles show swabs that were negative. (B) Whole genome sequences of four newly characterized white-tailed deer origin SARS-CoV-2 genomes were analyzed in the context of 135 publicly available white-tailed deer origin SARS-CoV-2 isolates and 63 arbitrarily selected SARS-CoV-2 Omicron genomes circulating amongst humans in New York City during this same time period as well as representative isolates from the environment or Syrian hamsters (SI Appendix, Table S2). The genome sequences were screened for quality, SNP positions called against the SARS-CoV-2 reference genome (NC_045512), and SNP alignments used to generate a maximum-likelihood phylogenetic tree using RAxML. The genome sequences from Staten Island white-tailed deer were genetically closely related to SARS-CoV-2 Omicron genomes recovered from humans in NY, as well as additional Omicron isolates from the environment.

To explore whether we might identify genetic changes associated with the observed “N” dropout in the TaqPath RT-PCR assay, several mutations were identified in the N gene, including D343G. However, the role, if any, of these mutations in the N gene in the TaqPath dropout remains to be determined since the primer and probe sequences for the CLIA approved TaqPath assay are proprietary and unavailable. Notably, the N dropout was only observed with the TaqPath assay and not in the OPTI Medical assay, which targets two separate regions in the N gene. N gene dropouts in RT-PCR assays have been reported with earlier SARS-CoV-2 VoCs^43,44^ and may therefore have the potential to not only affect assay performance but also might lead to false-negative results. Hence, further studies on the performance of current widely used RT-PCR assay are urgently warranted, since the occurrence of both S and N dropouts or amplification failures may compromise detection of continually emerging variants that are increasingly being observed in clinical samples.

### Is there evidence for SARS-CoV-2 re-infection of white-tailed deer?

An open question remains regarding whether deer, as in humans, may be re-infected with SARS-CoV-2 even in the presence of circulating neutralizing antibodies. Experimental SARS-CoV-2 infection in deer elicits a rapid neutralizing antibody immune response^9^, but it is not yet clear whether these antibodies prevent reinfection of deer. While direct evidence that experimentally or naturally SARS-CoV-2 infected white-tailed deer can be re-infected is lacking, it is noteworthy from the current investigation that one yearling male (number 2089) that was RT-PCR positive had a relatively robust level of neutralizing antibodies (78.7% inhibition, Table 1). It is possible that this deer may have rapidly seroconverted in response to SARS-CoV-2 infection even while continuing to shed viral RNA in nasal secretions as is observed in some SARS-CoV-2-positive human patients^45^. Alternatively, this animal may have seroconverted following a prior exposure to virus but remained susceptible to re-infection, as has also been observed to occur in humans^46^. These hypotheses need to be tested through larger field studies or through experimental infection investigations since the ability of the virus to overcome neutralizing antibody responses has significant consequences for our understanding of the overall transmission dynamics of SARS-CoV-2 in free-living deer and their potential for establishment as a reservoir host.

### Study Limitations

The sample size in our study is small in adult deer with respect to effect size and observed variation, and the samples come predominantly from young male deer. Hence, the inferences that might be derived for the larger population are limited. The study is also a cross-sectional survey of opportunistically sampled white-tailed deer in highly urbanized settings during an epizootic, with potentially greater opportunities for contact with humans than deer in other environments. Hence, the findings may not be broadly representative outside of these settings. Finally, the current analyses and sampling strategy do not enable us to differentiate whether animals positive for both viral RNA and neutralizing antibodies are newly seroconverted or individuals re-infected. This will require detailed laboratory investigations or longitudinal studies.

### Study implications and future directions

Given the well-documented susceptibility of white-tailed deer to SARS-CoV-2 as observed with experimental and natural infections, including with different viral lineages and VoCs, the current finding of spillover of Omicron is not altogether surprising. Previous studies have suggested that white-tailed deer remain largely asymptomatic following experimental SARS-CoV-2 infection^8,9^. However, we note that these trials utilized ancestral SARS-CoV-2 lineages including the B.1 and Alpha VoCs, and outcomes of infection of white-tailed deer infected with more recent SARS-CoV-2 variants, such as Omicron, are yet to be determined and should be the subject of future investigations. For instance, it is not known whether Omicron is more transmissible than Delta or the ancestral lineages in white-tailed deer as has been reported in humans. It is also unknown whether deer infection with Omicron results in clinical symptoms, unlike the experimental infections with ancestral SARS-CoV-2 strains and Alpha VoC that are primarily asymptomatic^8,9^.

Our finding of age-structure in sero-reactivity underscores the importance of better understanding social and behavioral drivers of transmission in free-living susceptible hosts and may have important implications for the development of future surveillance strategies. The ability of white-tailed deer to be re-infected with SARS-CoV-2, including with novel VoCs is also unknown. Our observation of a single individual positive by both the RT-PCR and sVNT assays is consistent with this possibility, though with our current methodology, it is not possible to determine the exact mechanisms by which this might occur. It will be important to determine through future investigations whether white-tailed deer have durable immunity or are susceptible to re-infection to better model long-term trajectories. Such additional information is important to inform the parameterization of disease dynamic models to predict likelihood and conditions needed for the virus becoming endemic or fading out in free-living deer populations.

It is well established that recombination and adaptation events contribute to new coronavirus species, enabling successful host shifts and jumps^47^. The lower observed concordance of N protein serological reactivity with sVNT (Cohen’s Kappa 0.32) as compared with RBD and sVNT (Cohen’s Kappa 0.92) together with the considerable number of individuals that had antibodies to N protein but not to either RBD (12 of 23) or sVNT (14 of 23) (Supplemental Table 1), suggests that white-tailed deer on Staten Island appear likely to have been exposed to endemic coronaviruses. Combined with the previously observed high viral titers reported in deer retropharyngeal lymph nodes and palatine tonsils^6^, this suggests that opportunities for recombination between SARS-CoV-2 and deer coronaviruses may occur; such scenarios may have important consequence for the emergence of new variants and deserves urgent further investigation.

The multiple human-to-animal spillovers and documentation of subsequent onward transmission in free-living mink and deer also highlight the ‘generalist’ and rapidly adaptive nature of SARS-CoV-2 as a pathogen of mammalian hosts as recently described^48^. Emerging evidence suggests host adaption of SARS-Cov-2 in animal hosts and six specific mutations in mink (NSP9_G37E, Spike_F486L, Spike_N501T, Spike_Y453F, ORF3a_T229I, ORF3a_L219V), and one in deer (NSP3a_L1035F) have been identified^48^. Given the importance of monitoring for potential for spillback to humans, our findings highlight the importance of longitudinal studies to monitor SARS-CoV-2 adaptation and evolution within free-living deer as well as in other susceptible non-human animal populations.

Several wild and peri-domestic animal species susceptible to SARS-CoV-2 infection share the same ecological space with deer. However, widespread SARS-CoV-2 spillovers into wild animals other than to white-tailed deer (and perhaps mink) are currently unknown. While SARS-CoV-2 infections in white-tailed deer have reported in 15 States in North America^10^, infection of captive or free-living cervids across different parts of the world remains completely unknown and also needs investigation.

Finally, given the expanding reported host-range of species susceptible to SARS-CoV-2 and the generalist nature of the virus^3,48^, there is an urgent unmet need for well-coordinated targeted surveillance of at-risk wild animal species, including white-tailed deer, to better understand the transmission networks and assess risk of spillback to humans.

## Acknowledgements

We thank, Lindsey LaBella, Shubhada K. Chothe, Padmaja Jakka, Abirami Ravichandran and Nishitha Kodali for help with PCR and sample processing. We thank Matthew Ojeda Saavedra, Sindy Pena, Kristina Reppond, Madison N. Shyer, Jessica Cambric, Ryan Gadd, Rashi M. Thakur, Akanksha Batajoo, and Regan Mangham for genome sequencing. Samples used in this study came from deer handled under authorization provided through license issued by New York State Department of Environmental Conservation. We are grateful to Drs. Suelee Robbe-Austerman, Kristina Lantz, Mia Kim Torchetti, and Julianna Lenoch of the USDA-APHIS for their invaluable assistance and wise counsel during the investigation. We thank Dr. Gleyder Roman-Sosa for the recombinant S and N antigen preparation. This study was made possible by the collective efforts of White Buffalo, Inc., the City of New York Parks & Recreation and their field staff for help with sample collection.

The study was funded by the Huck institutes of the Life Sciences (S.V.K. and V.K.), US Department of Agriculture National Institute of Food and Agriculture (NIFA) Award 2020-67015-32175. This project was also funded in part by the Houston Methodist Academic Institute Infectious Diseases Fund (J.M.M. and R.J.O.) and supported in part by National Institute of Allergy and Infectious Diseases, National Institutes of Health, Department of Health and Human Services, Contract No. 75N93019C00076 (J.J.D.). K.V. was partially supported by the NSF Ecology and Evolution of Infectious Diseases program (Grant #1619072) as well as a grant from the NIH (1R21AI156406-01). J.A.R was partially funded by the USDA-NIFA Program under award number 2020-67015-33157, the AMP Core of the Center of Emerging and Zoonotic Infectious Diseases (CEZID) from National Institute of General Medical Sciences (NIGMS) under award number P20GM130448, and the NIAID Centers of Excellence for Influenza Research and Surveillance under contract number HHSN 272201400006C.

