## Supplemental Table 1 for "Detection of SARS-CoV-2 Omicron variant (B.1.1.529) infection of white-tailed deer"

**Supplementary Table 1. Metadata associated with the deer samples collected from Staten Island, NY**

| ID | Capture Date | Age (years) | Sex | sVN result | sVN (inhibition %) | RBD_ELISA results | RBD_ELISA OD | N_ELISA results | N_ELISA OD | Rt-PCR results | Rt-PCR Ct |
| --- | --- | --- | --- | --- | --- | --- | --- | --- | --- | --- | --- |
| 80 | 12/12/2021 | 0.5 | M | NEG | 3.73 | NEG | 0.14 | NEG | 0.26 | ND | ND |
| 90 | 1/11/2022 | 0.5 | F | NEG | 10.58 | NEG | 0.14 | NEG | 0.29 | NEG | UN |
| 97 | 1/19/2022 | 0.5 | F | NEG | 7.44 | ND | ND | ND | ND | ND | ND |
| 100 | 1/9/2022 | 0.5 | F | NEG | 8.89 | NEG | 0.14 | NEG | 0.31 | NEG | UN |
| 101 | 1/20/2022 | 3.5 | F | NEG | 7.71 | ND | ND | ND | ND | NEG | UN |
| 102 | 1/6/2022 | 0.5 | F | ND | ND | NEG | 0.14 | NEG | 0.32 | POS | 30.64 |
| 103 | 1/6/2022 | 0.5 | F | NEG | 8.85 | NEG | 0.14 | NEG | 0.34 | POS | 32.14 |
| 104 | 1/6/2022 | 0.5 | F | NEG | 7.81 | NEG | 0.14 | NEG | 0.27 | POS | 35.57 |
| 106 | 1/27/2022 | 2.5 | F | NEG | 22.69 | ND | ND | ND | ND | NEG | UN |
| 120 | 12/18/2021 | 0.5 | F | NEG | 10.99 | NEG | 0.14 | NEG | 0.34 | ND | ND |
| 121 | 1/9/2022 | 0.5 | F | NEG | 11.31 | NEG | 0.14 | NEG | 0.34 | ND | ND |
| 123 | 1/5/2022 | 0.5 | F | NEG | 2.56 | NEG | 0.14 | NEG | 0.31 | NEG | UN |
| 1584 | 12/12/2021 | 3.5 | M | POS | 74.72 | POS | 0.21 | POS | 0.59 | ND | ND |
| 2002 | 12/15/2021 | 1.5 | M | NEG | 28.82 | POS | 0.17 | POS | 0.44 | ND | ND |
| 2003 | 12/14/2021 | 2.5 | M | NEG | 2.50 | NEG | 0.14 | NEG | 0.38 | ND | ND |
| 2004 | 12/13/2021 | 1.5 | M | POS | 93.97 | POS | 0.29 | POS | 0.48 | ND | ND |
| 2005 | 12/13/2021 | 0.5 | M | NEG | 8.40 | NEG | 0.16 | NEG | 0.38 | ND | ND |
| 2006 | 12/29/2021 | 0.5 | M | NEG | 2.50 | NEG | 0.13 | NEG | 0.22 | ND | ND |
| 2007 | 12/29/2021 | 0.5 | M | NEG | 3.08 | NEG | 0.14 | NEG | 0.26 | ND | ND |
| 2008 | 12/15/2021 | 1.5 | M | NEG | 0.36 | NEG | 0.15 | POS | 0.43 | ND | ND |
| 2009 | 12/15/2021 | 1.5 | M | NEG | -2.11 | NEG | 0.14 | NEG | 0.31 | ND | ND |
| 2010 | 12/15/2021 | 0.5 | M | NEG | 2.82 | NEG | 0.14 | NEG | 0.26 | ND | ND |
| 2011 | 12/16/2021 | 1.5 | M | NEG | 3.01 | NEG | 0.14 | NEG | 0.37 | ND | ND |
| 2012 | 12/16/2021 | 0.5 | M | NEG | 3.27 | NEG | 0.15 | NEG | 0.37 | ND | ND |
| 2013 | 12/15/2021 | 0.5 | M | NEG | 3.47 | NEG | 0.15 | NEG | 0.22 | ND | ND |
| 2014 | 12/20/2021 | 1.5 | M | NEG | 4.51 | NEG | 0.14 | POS | 0.71 | ND | ND |
| 2015 | 12/17/2021 | 0.5 | M | POS | 87.75 | POS | 0.25 | NEG | 0.37 | ND | ND |
| 2016 | 12/18/2021 | 0.5 | M | NEG | -6.84 | NEG | 0.15 | NEG | 0.31 | ND | ND |
| 2018 | 12/18/2021 | 1.5 | M | NEG | 9.82 | NEG | 0.14 | NEG | 0.32 | ND | ND |
| 2019 | 12/19/2021 | 2.5 | M | NEG | 8.14 | NEG | 0.15 | POS | 0.43 | ND | ND |
| 2020 | 12/16/2021 | 0.5 | M | NEG | 2.30 | NEG | 0.14 | NEG | 0.29 | ND | ND |
| 2021 | 12/19/2021 | 0.5 | M | NEG | 0.75 | NEG | 0.14 | NEG | 0.26 | ND | ND |
| 2022 | 12/17/2021 | 0.5 | M | NEG | 5.28 | NEG | 0.14 | NEG | 0.33 | ND | ND |
| 2023 | 12/22/2021 | 1.5 | M | NEG | 3.92 | NEG | 0.15 | NEG | 0.36 | ND | ND |
| 2025 | 12/18/2021 | 1.5 | M | POS | 92.35 | POS | 0.42 | POS | 0.47 | ND | ND |
| 2026 | 12/18/2021 | 0.5 | M | NEG | 3.73 | NEG | 0.14 | NEG | 0.24 | ND | ND |
| 2028 | 12/20/2021 | 0.5 | M | NEG | 6.77 | NEG | 0.14 | NEG | 0.31 | ND | ND |
| 2029 | 12/20/2021 | 0.5 | M | NEG | 1.52 | NEG | 0.14 | NEG | 0.22 | ND | ND |
| 2031 | 12/22/2021 | 0.5 | M | NEG | 2.50 | NEG | 0.14 | NEG | 0.27 | ND | ND |
| ID | **Capture Date** | **Age (years)** | **Sex** | **sVN result** | **sVN (inhibition %)** | **RBD_ELISA results** | **RBD_ELISA OD** | **N_ELISA results** | **N_ELISA OD** | **Rt-PCR results** | **Rt-PCR Ct** |
| 2032 | 12/21/2021 | 0.5 | M | NEG | 2.24 | NEG | 0.14 | NEG | 0.25 | ND | ND |
| 2033 | 12/31/2021 | 1.5 | M | POS | 68.62 | POS | 0.2 | NEG | 0.29 | ND | ND |
| 2034 | 12/30/2021 | 0.5 | M | NEG | 3.21 | NEG | 0.14 | NEG | 0.29 | ND | ND |
| 2035 | 12/21/2021 | 1.5 | M | POS | 33.16 | POS | 0.17 | POS | 0.45 | ND | ND |
| 2036 | 12/28/2021 | 1.5 | M | POS | 60.00 | POS | 0.18 | NEG | 0.38 | ND | ND |
| 2038 | 12/22/2021 | 0.5 | M | NEG | 15.59 | POS | 0.19 | POS | 0.5 | ND | ND |
| 2039 | 12/29/2021 | 0.5 | M | NEG | 19.55 | NEG | 0.14 | NEG | 0.25 | ND | ND |
| 2040 | 12/30/2021 | 1.5 | M | POS | 51.83 | POS | 0.19 | NEG | 0.33 | ND | ND |
| 2041 | 12/30/2021 | 1.5 | M | NEG | 10.08 | NEG | 0.14 | NEG | 0.38 | ND | ND |
| 2042 | 12/31/2021 | 0.5 | M | NEG | 11.31 | NEG | 0.14 | NEG | 0.38 | ND | ND |
| 2043 | 1/1/2022 | 0.5 | M | NEG | 2.50 | NEG | 0.14 | NEG | 0.29 | ND | ND |
| 2044 | 12/31/2021 | 1.5 | M | NEG | 1.07 | NEG | 0.15 | POS | 0.56 | ND | ND |
| 2045 | 12/30/2021 | 1.5 | M | POS | 46.97 | POS | 0.19 | POS | 0.48 | ND | ND |
| 2047 | 1/2/2022 | 2.5 | M | NEG | 16.24 | NEG | 0.15 | NEG | 0.37 | ND | ND |
| 2048 | 1/3/2022 | 0.5 | M | NEG | 10.47 | NEG | 0.13 | NEG | 0.24 | NEG | UN |
| 2049 | 1/4/2022 | 1.5 | M | NEG | 15.82 | NEG | 0.14 | POS | 0.5 | NEG | UN |
| 2050 | 12/31/2021 | 1.5 | M | NEG | 3.47 | NEG | 0.14 | POS | 0.51 | ND | ND |
| 2051 | 1/3/2022 | 1.5 | M | NEG | 11.31 | NEG | 0.14 | NEG | 0.3 | NEG | UN |
| 2052 | 1/1/2022 | 1.5 | M | NEG | 1.85 | NEG | 0.14 | NEG | 0.28 | ND | ND |
| 2053 | 1/4/2022 | 1.5 | M | NEG | 7.23 | NEG | 0.14 | NEG | 0.29 | NEG | UN |
| 2054 | 1/4/2022 | 0.5 | M | NEG | 0.94 | NEG | 0.14 | NEG | 0.24 | NEG | UN |
| 2055 | 1/4/2022 | 1.5 | M | POS | 97.02 | POS | 0.62 | NEG | 0.34 | NEG | UN |
| 2056 | 1/7/2022 | 0.5 | M | NEG | 3.29 | NEG | 0.15 | NEG | 0.37 | NEG | UN |
| 2057 | 1/2/2022 | 0.5 | M | NEG | 6.06 | NEG | 0.14 | NEG | 0.25 | ND | ND |
| 2058 | 1/2/2022 | 0.5 | M | NEG | 8.14 | NEG | 0.13 | NEG | 0.26 | ND | ND |
| 2059 | 1/7/2022 | 3.5 | M | NEG | 4.64 | NEG | 0.14 | POS | 0.4 | NEG | UN |
| 2060 | 1/2/2022 | 1.5 | M | NEG | 6.04 | NEG | 0.15 | POS | 0.44 | ND | ND |
| 2061 | 1/8/2022 | 0.5 | M | NEG | 7.82 | NEG | 0.14 | NEG | 0.32 | NEG | UN |
| 2062 | 1/5/2022 | 1.5 | M | POS | 70.89 | POS | 0.2 | POS | 0.49 | NEG | UN |
| 2063 | 1/6/2022 | 1.5 | M | ND | ND | ND | ND | ND | ND | NEG | UN |
| 2064 | 1/5/2022 | 0.5 | M | POS | 69.66 | POS | 0.22 | NEG | 0.36 | NEG | UN |
| 2065 | 1/6/2022 | 0.5 | M | ND | ND | ND | ND | ND | ND | NEG | UN |
| 2066 | 1/7/2022 | 1.5 | M | NEG | -0.80 | NEG | 0.14 | NEG | 0.33 | NEG | UN |
| 2067 | 1/7/2022 | 0.5 | M | NEG | 7.36 | NEG | 0.13 | NEG | 0.24 | POS | 36.3 |
| 2068 | 1/9/2022 | 0.5 | M | ND | ND | ND | ND | ND | ND | NEG | UN |
| 2070 | 1/6/2022 | 1.5 | M | NEG | 8.91 | NEG | 0.14 | NEG | 0.32 | ND | ND |
| 2071 | 1/8/2022 | 2.5 | M | POS | 96.37 | POS | 0.35 | POS | 0.54 | NEG | UN |
| 2072 | 1/13/2022 | 0.5 | M | NEG | 14.42 | NEG | 0.14 | NEG | 0.3 | NEG | UN |
| 2073 | 1/10/2022 | 0.5 | M | NEG | 12.80 | NEG | 0.14 | NEG | 0.27 | NEG | UN |
| 2074 | 1/8/2022 | 0.5 | M | NEG | 7.47 | NEG | 0.15 | NEG | 0.34 | NEG | UN |
| 2075 | 1/14/2022 | 1.5 | M | POS | 78.48 | POS | 0.22 | NEG | 0.34 | NEG | UN |
| ID | **Capture Date** | **Age (years)** | **Sex** | **sVN result** | **sVN (inhibition %)** | **RBD_ELISA results** | **RBD_ELISA OD** | **N_ELISA results** | **N_ELISA OD** | **Rt-PCR results** | **Rt-PCR Ct** |
| 2076 | 1/9/2022 | 0.5 | M | NEG | 11.90 | NEG | 0.14 | NEG | 0.29 | NEG | UN |
| 2077 | 1/10/2022 | 1.5 | M | POS | 34.31 | POS | 0.18 | POS | 0.39 | NEG | UN |
| 2078 | 1/10/2022 | 0.5 | M | NEG | 5.22 | NEG | 0.14 | NEG | 0.3 | NEG | UN |
| 2079 | 1/10/2022 | 2.5 | M | NEG | 6.84 | NEG | 0.14 | POS | 0.4 | NEG | UN |
| 2080 | 1/9/2022 | 0.5 | M | NEG | 20.52 | NEG | 0.14 | POS | 0.39 | NEG | UN |
| 2081 | 1/12/2022 | 2.5 | M | NEG | 10.41 | NEG | 0.14 | NEG | 0.32 | NEG | UN |
| 2082 | 1/10/2022 | 1.5 | M | NEG | 10.28 | NEG | 0.14 | NEG | 0.29 | NEG | UN |
| 2083 | 1/11/2022 | 0.5 | M | NEG | 5.87 | NEG | 0.13 | NEG | 0.26 | NEG | UN |
| 2084 | 1/10/2022 | 0.5 | M | NEG | 10.28 | NEG | 0.14 | NEG | 0.37 | NEG | UN |
| 2085 | 1/12/2022 | 3.5 | M | NEG | 3.73 | NEG | 0.14 | NEG | 0.3 | NEG | UN |
| 2086 | 1/11/2022 | 1.5 | M | NEG | 12.22 | NEG | 0.14 | POS | 0.48 | NEG | UN |
| 2087 | 1/15/2022 | 0.5 | M | NEG | 0.09 | NEG | 0.14 | NEG | 0.2 | NEG | UN |
| 2088 | 1/14/2022 | 2.5 | M | NEG | 9.91 | NEG | 0.14 | POS | 0.49 | NEG | UN |
| 2089 | 1/16/2022 | 1.5 | M | POS | 78.74 | POS | 0.24 | POS | 0.52 | POS | 36.12 |
| 2090 | 1/15/2022 | 0.5 | M | NEG | 8.98 | NEG | 0.14 | NEG | 0.23 | NEG | UN |
| 2091 | 1/11/2022 | 2.5 | M | NEG | 0.94 | NEG | 0.16 | NEG | 0.3 | NEG | UN |
| 2092 | 1/25/2022 | 0.5 | M | NEG | -5.83 | ND | ND | ND | ND | ND | ND |
| 2093 | 1/16/2022 | 0.5 | M | NEG | 6.19 | NEG | 0.14 | NEG | 0.37 | NEG | UN |
| 2094 | 1/14/2022 | 0.5 | M | ND | ND | ND | ND | ND | ND | NEG | UN |
| 2095 | 1/20/2022 | 3.5 | M | NEG | 3.14 | ND | ND | ND | ND | NEG | UN |
| 2096 | 1/16/2022 | 0.5 | M | NEG | 6.00 | NEG | 0.14 | NEG | 0.29 | NEG | UN |
| 2097 | 1/15/2022 | 0.5 | M | NEG | 12.16 | NEG | 0.14 | NEG | 0.31 | NEG | UN |
| 2098 | 1/17/2022 | 0.5 | M | NEG | 5.20 | ND | ND | ND | ND | NEG | UN |
| 2099 | 1/12/2022 | 0.5 | M | NEG | 13.52 | NEG | 0.13 | NEG | 0.2 | NEG | UN |
| 2100 | 1/17/2022 | 0.5 | M | NEG | 9.87 | ND | ND | ND | ND | NEG | UN |
| 2101 | 1/23/2022 | 0.5 | M | NEG | 13.36 | ND | ND | ND | ND | ND | ND |
| 2102 | 1/13/2022 | 0.5 | M | NEG | 24.15 | NEG | 0.15 | NEG | 0.32 | NEG | UN |
| 2103 | 1/18/2022 | 0.5 | M | POS | 93.99 | ND | ND | ND | ND | NEG | UN |
| 2104 | 1/24/2022 | 0.5 | M | NEG | -3.05 | ND | ND | ND | ND | ND | ND |
| 2105 | 1/18/2022 | 0.5 | M | NEG | -3.41 | ND | ND | ND | ND | NEG | UN |
| 2106 | 1/20/2022 | 2.5 | M | NEG | 12.02 | ND | ND | ND | ND | NEG | UN |
| 2107 | 1/23/2022 | 0.5 | M | NEG | 3.05 | ND | ND | ND | ND | ND | ND |
| 2108 | 1/27/2022 | 0.5 | M | NEG | 5.11 | ND | ND | ND | ND | POS | 22.5 |
| 2109 | 1/21/2022 | 0.5 | M | POS | 83.14 | ND | ND | ND | ND | ND | ND |
| 2110 | 1/18/2022 | 0.5 | M | NEG | 3.95 | ND | ND | ND | ND | NEG | UN |
| 2111 | 1/24/2022 | 0.5 | M | NEG | 6.64 | ND | ND | ND | ND | ND | ND |
| 2112 | 1/25/2022 | 0.5 | M | NEG | -10.49 | ND | ND | ND | ND | ND | ND |
| 2113 | 1/24/2022 | 0.5 | M | NEG | -8.52 | ND | ND | ND | ND | ND | ND |
| 2114 | 1/25/2022 | 0.5 | M | NEG | 8.61 | ND | ND | ND | ND | ND | ND |
| 2115 | 1/26/2022 | 0.5 | M | NEG | 4.30 | ND | ND | ND | ND | ND | ND |
| 2116 | 1/25/2022 | 1.5 | M | NEG | 5.38 | ND | ND | ND | ND | ND | ND |
| ID | **Capture Date** | **Age (years)** | **Sex** | **sVN result** | **sVN (inhibition %)** | **RBD_ELISA results** | **RBD_ELISA OD** | **N_ELISA results** | **N_ELISA OD** | **Rt-PCR results** | **Rt-PCR Ct** |
| 2117 | 1/28/2022 | 0.5 | M | NEG | 6.64 | ND | ND | ND | ND | NEG | UN |
| 2118 | 1/24/2022 | 0.5 | M | NEG | 2.42 | ND | ND | ND | ND | ND | ND |
| 2119 | 1/26/2022 | 0.5 | M | NEG | 8.79 | ND | ND | ND | ND | ND | ND |
| 2120 | 1/27/2022 | 0.5 | M | NEG | 1.08 | ND | ND | ND | ND | NEG | UN |
| 2121 | 1/25/2022 | 0.5 | M | NEG | 6.19 | ND | ND | ND | ND | ND | ND |
| 2122 | 1/26/2022 | 0.5 | M | NEG | 2.60 | ND | ND | ND | ND | ND | ND |
| 2123 | 1/27/2022 | 2.5 | M | NEG | 24.66 | ND | ND | ND | ND | NEG | UN |
| 2124 | 1/27/2022 | 1.5 | M | POS | 92.38 | ND | ND | ND | ND | NEG | UN |
| 2125 | 1/30/2022 | 0.5 | M | NEG | 4.13 | ND | ND | ND | ND | POS | 31.1 |
| 2126 | 1/30/2022 | 0.5 | M | NEG | 1.79 | ND | ND | ND | ND | NEG | UN |
| 40101 | 1/19/2022 | 0.5 | F | NEG | -0.27 | ND | ND | ND | ND | NEG | UN |
| 40105 | 1/19/2022 | 0.5 | F | NEG | 2.24 | ND | ND | ND | ND | NEG | UN |
| 40109 | 1/18/2022 | 0.5 | F | NEG | -13.18 | ND | ND | ND | ND | ND | ND |
| 40115 | 1/23/2022 | 0.5 | F | NEG | 11.75 | ND | ND | ND | ND | ND | ND |
| 40129 | 1/20/2022 | 0.5 | F | NEG | 8.34 | ND | ND | ND | ND | NEG | UN |

\# F: Female; M: Male; sVN: Virus neutralization using the sVNT assay; NEG: Negative; POS: Positive; ND: Not determined; RBD_ELISA, Receptor Binding Domain enzyme-linked immunosorbent assay; N_ELISA: Nucleocapsid enzyme-linked immunosorbent assay; OD: optical density; UN: Unknown
