## Supplemental Table 2 for "Detection of SARS-CoV-2 Omicron variant (B.1.1.529) infection of white-tailed deer"

**Supplemental Table 2.** Metadata associated with the taxa represented in the phylogenetic tree presented in Figure 2.*

| Tree node ID | Host | Collection date | Lineage | Location |
| --- | --- | --- | --- | --- |
| NY WTD102 | White-Tailed Deer | 1/6/2022 | BA.1 | New York, USA |
| NY WTD103 | White-Tailed Deer | 1/6/2022 | BA.1 | New York, USA |
| NY WTD104 | White-Tailed Deer | 1/6/2022 | BA.1 | New York, USA |
| NY WTD2067 | White-Tailed Deer | 1/7/2022 | BA.1 | New York, USA |
| hCoV-19/deer/USA/AR-21-038764-001/2021 | White-Tailed Deer | 12/1/2021 | AY.54 | Arkansas, USA |
| hCoV-19/deer/USA/AR-21-038764-002/2021 | White-Tailed Deer | 12/1/2021 | AY.45 | Arkansas, USA |
| hCoV-19/deer/USA/IA-200069/2020 | White-Tailed Deer | 12/5/2020 | B.1.119 | Iowa, USA |
| hCoV-19/deer/USA/IA-200191/2020 | White-Tailed Deer | 12/8/2020 | B.1.311 | Iowa, USA |
| hCoV-19/deer/USA/IA-200192/2020 | White-Tailed Deer | 12/8/2020 | B.1.119 | Iowa, USA |
| hCoV-19/deer/USA/IA-200193/2020 | White-Tailed Deer | 12/8/2020 | B.1.311 | Iowa, USA |
| hCoV-19/deer/USA/IA-200194/2020 | White-Tailed Deer | 12/8/2020 | B.1.311 | Iowa, USA |
| hCoV-19/deer/USA/IA-200195/2020 | White-Tailed Deer | 12/8/2020 | B.1.311 | Iowa, USA |
| hCoV-19/deer/USA/IA-200239/2020 | White-Tailed Deer | 12/5/2020 | B.1.264 | Iowa, USA |
| hCoV-19/deer/USA/IA-200442/2021 | White-Tailed Deer | 1/9/2021 | B.1.311 | Iowa, USA |
| hCoV-19/deer/USA/IA-200443/2021 | White-Tailed Deer | 1/9/2021 | B.1.311 | Iowa, USA |
| hCoV-19/deer/USA/IA-200444/2021 | White-Tailed Deer | 1/9/2021 | B.1.311 | Iowa, USA |
| hCoV-19/deer/USA/IA-200566/2020 | White-Tailed Deer | 12/5/2020 | B.1.311 | Iowa, USA |
| hCoV-19/deer/USA/IA-200567/2020 | White-Tailed Deer | 12/5/2020 | B.1.311 | Iowa, USA |
| hCoV-19/deer/USA/IA-200568/2020 | White-Tailed Deer | 12/5/2020 | B.1.311 | Iowa, USA |
| hCoV-19/deer/USA/IA-200569/2020 | White-Tailed Deer | 12/5/2020 | B.1.311 | Iowa, USA |
| hCoV-19/deer/USA/IA-200570/2020 | White-Tailed Deer | 12/5/2020 | B.1.311 | Iowa, USA |
| hCoV-19/deer/USA/IA-200571/2020 | White-Tailed Deer | 12/5/2020 | B.1.311 | Iowa, USA |
| hCoV-19/deer/USA/IA-200572/2020 | White-Tailed Deer | 12/5/2020 | B.1.311 | Iowa, USA |
| hCoV-19/deer/USA/IA-200573/2020 | White-Tailed Deer | 12/5/2020 | B.1.311 | Iowa, USA |
| hCoV-19/deer/USA/IA-200574/2020 | White-Tailed Deer | 12/5/2020 | B.1.311 | Iowa, USA |
| hCoV-19/deer/USA/IA-200772/2020 | White-Tailed Deer | 12/5/2020 | B.1.311 | Iowa, USA |
| Tree node ID | **Host** | **Collection date** | **Lineage** | **Location** |
| hCoV-19/deer/USA/IA-200862/2021 | White-Tailed Deer | 1/9/2021 | B.1.311 | Iowa, USA |
| hCoV-19/deer/USA/IA-200866/2021 | White-Tailed Deer | 1/9/2021 | B.1.311 | Iowa, USA |
| hCoV-19/deer/USA/IA-201788/2020 | White-Tailed Deer | 12/6/2020 | B.1 | Iowa, USA |
| hCoV-19/deer/USA/IA-201793/2020 | White-Tailed Deer | 12/6/2020 | B.1.240 | Iowa, USA |
| hCoV-19/deer/USA/IA-201794/2020 | White-Tailed Deer | 12/6/2020 | B.1.311 | Iowa, USA |
| hCoV-19/deer/USA/IA-201795/2020 | White-Tailed Deer | 12/6/2020 | B.1 | Iowa, USA |
| hCoV-19/deer/USA/IA-201796/2020 | White-Tailed Deer | 12/6/2020 | B.1.240 | Iowa, USA |
| hCoV-19/deer/USA/IA-201797/2020 | White-Tailed Deer | 12/6/2020 | B.1 | Iowa, USA |
| hCoV-19/deer/USA/IA-201820/2020 | White-Tailed Deer | 12/10/2020 | B.1.234 | Iowa, USA |
| hCoV-19/deer/USA/IA-201821/2020 | White-Tailed Deer | 12/10/2020 | B.1.234 | Iowa, USA |
| hCoV-19/deer/USA/IA-201833/2020 | White-Tailed Deer | 12/10/2020 | B.1.234 | Iowa, USA |
| hCoV-19/deer/USA/IA-201835/2020 | White-Tailed Deer | 12/10/2020 | B.1.234 | Iowa, USA |
| hCoV-19/deer/USA/IA-203522/2020 | White-Tailed Deer | 12/8/2020 | B.1 | Iowa, USA |
| hCoV-19/deer/USA/IA-203525/2020 | White-Tailed Deer | 12/8/2020 | B.1 | Iowa, USA |
| hCoV-19/deer/USA/IA-203526/2020 | White-Tailed Deer | 12/8/2020 | B.1 | Iowa, USA |
| hCoV-19/deer/USA/IA-203700/2020 | White-Tailed Deer | 12/9/2020 | B.1 | Iowa, USA |
| hCoV-19/deer/USA/IA-203701/2020 | White-Tailed Deer | 12/9/2020 | B.1 | Iowa, USA |
| hCoV-19/deer/USA/IA-203704/2020 | White-Tailed Deer | 12/9/2020 | B.1.400 | Iowa, USA |
| hCoV-19/deer/USA/IA-203705/2020 | White-Tailed Deer | 12/9/2020 | B.1.2 | Iowa, USA |
| hCoV-19/deer/USA/IA-204363/2020 | White-Tailed Deer | 12/7/2020 | B.1 | Iowa, USA |
| hCoV-19/deer/USA/IA-204364/2020 | White-Tailed Deer | 12/7/2020 | B.1.2 | Iowa, USA |
| hCoV-19/deer/USA/IA-204472/2020 | White-Tailed Deer | 12/5/2020 | B.1.2 | Iowa, USA |
| hCoV-19/deer/USA/IA-204473/2020 | White-Tailed Deer | 12/5/2020 | B.1.2 | Iowa, USA |
| hCoV-19/deer/USA/IA-204478/2020 | White-Tailed Deer | 12/5/2020 | B.1 | Iowa, USA |
| hCoV-19/deer/USA/IA-204668/2020 | White-Tailed Deer | 12/9/2020 | B.1 | Iowa, USA |
| hCoV-19/deer/USA/IA-204669/2020 | White-Tailed Deer | 12/9/2020 | B.1.2 | Iowa, USA |
| hCoV-19/deer/USA/IA-205750/2020 | White-Tailed Deer | 12/7/2020 | B.1.2 | Iowa, USA |
| Tree node ID | **Host** | **Collection date** | **Lineage** | **Location** |
| hCoV-19/deer/USA/IA-205769/2020 | White-Tailed Deer | 12/8/2020 | B.1.2 | Iowa, USA |
| hCoV-19/deer/USA/IA-205795/2020 | White-Tailed Deer | 12/7/2020 | B.1.2 | Iowa, USA |
| hCoV-19/deer/USA/IA-206407/2020 | White-Tailed Deer | 11/6/2020 | B.1.2 | Iowa, USA |
| hCoV-19/deer/USA/IA-206527/2020 | White-Tailed Deer | 10/15/2020 | B.1.2 | Iowa, USA |
| hCoV-19/deer/USA/IA-206535/2020 | White-Tailed Deer | 12/22/2020 | B.1.234 | Iowa, USA |
| hCoV-19/deer/USA/IA-206739/2020 | White-Tailed Deer | 12/15/2020 | B.1.234 | Iowa, USA |
| hCoV-19/deer/USA/IA-206978/2020 | White-Tailed Deer | 12/28/2020 | B.1.2 | Iowa, USA |
| hCoV-19/deer/USA/IA-207150/2020 | White-Tailed Deer | 11/15/2020 | B.1.2 | Iowa, USA |
| hCoV-19/deer/USA/IA-207253/2020 | White-Tailed Deer | 11/30/2020 | B.1.2 | Iowa, USA |
| hCoV-19/deer/USA/IA-207337/2020 | White-Tailed Deer | 9/30/2020 | B.1.2 | Iowa, USA |
| hCoV-19/deer/USA/IA-207391/2020 | White-Tailed Deer | 11/29/2020 | B.1.2 | Iowa, USA |
| hCoV-19/deer/USA/IA-207431/2020 | White-Tailed Deer | 10/28/2020 | B.1.2 | Iowa, USA |
| hCoV-19/deer/USA/IA-207437/2020 | White-Tailed Deer | 10/31/2020 | B.1.2 | Iowa, USA |
| hCoV-19/deer/USA/IA-207530/2020 | White-Tailed Deer | 11/7/2020 | B.1.2 | Iowa, USA |
| hCoV-19/deer/USA/IA-207531/2020 | White-Tailed Deer | 11/8/2020 | B.1.2 | Iowa, USA |
| hCoV-19/deer/USA/IA-207579/2020 | White-Tailed Deer | 11/17/2020 | B.1.2 | Iowa, USA |
| hCoV-19/deer/USA/IA-207589/2020 | White-Tailed Deer | 11/24/2020 | B.1.2 | Iowa, USA |
| hCoV-19/deer/USA/IA-207649/2020 | White-Tailed Deer | 12/5/2020 | B.1.2 | Iowa, USA |
| hCoV-19/deer/USA/IA-207707/2020 | White-Tailed Deer | 12/10/2020 | B.1.2 | Iowa, USA |
| hCoV-19/deer/USA/IA-207770/2020 | White-Tailed Deer | 12/2/2020 | B.1.2 | Iowa, USA |
| hCoV-19/deer/USA/IA-208017/2020 | White-Tailed Deer | 9/28/2020 | B.1.2 | Iowa, USA |
| hCoV-19/deer/USA/IA-208043/2020 | White-Tailed Deer | 10/8/2020 | B.1.2 | Iowa, USA |
| hCoV-19/deer/USA/IA-208095/2020 | White-Tailed Deer | 11/24/2020 | B.1.2 | Iowa, USA |
| hCoV-19/deer/USA/IA-208098/2020 | White-Tailed Deer | 11/24/2020 | B.1.2 | Iowa, USA |
| hCoV-19/deer/USA/IA-208099/2020 | White-Tailed Deer | 11/24/2020 | B.1.2 | Iowa, USA |
| hCoV-19/deer/USA/IA-208100/2020 | White-Tailed Deer | 11/24/2020 | B.1.2 | Iowa, USA |
| hCoV-19/deer/USA/IA-208101/2020 | White-Tailed Deer | 11/24/2020 | B.1.2 | Iowa, USA |
| Tree node ID | **Host** | **Collection date** | **Lineage** | **Location** |
| hCoV-19/deer/USA/IA-208102/2020 | White-Tailed Deer | 11/24/2020 | B.1.2 | Iowa, USA |
| hCoV-19/deer/USA/IA-208103/2020 | White-Tailed Deer | 11/24/2020 | B.1.2 | Iowa, USA |
| hCoV-19/deer/USA/IA-208104/2020 | White-Tailed Deer | 11/24/2020 | B.1.2 | Iowa, USA |
| hCoV-19/deer/USA/IA-208105/2020 | White-Tailed Deer | 11/24/2020 | B.1.2 | Iowa, USA |
| hCoV-19/deer/USA/IA-208106/2020 | White-Tailed Deer | 11/24/2020 | B.1.2 | Iowa, USA |
| hCoV-19/deer/USA/IA-208107/2020 | White-Tailed Deer | 11/24/2020 | B.1.2 | Iowa, USA |
| hCoV-19/deer/USA/IA-208108/2020 | White-Tailed Deer | 12/10/2020 | B.1.2 | Iowa, USA |
| hCoV-19/deer/USA/IA-208109/2020 | White-Tailed Deer | 12/10/2020 | B.1.2 | Iowa, USA |
| hCoV-19/deer/USA/IA-208110/2020 | White-Tailed Deer | 12/10/2020 | B.1.2 | Iowa, USA |
| hCoV-19/deer/USA/IA-208111/2020 | White-Tailed Deer | 12/10/2020 | B.1.2 | Iowa, USA |
| hCoV-19/deer/USA/IA-208112/2020 | White-Tailed Deer | 12/10/2020 | B.1.2 | Iowa, USA |
| hCoV-19/deer/USA/IA-208113/2020 | White-Tailed Deer | 12/10/2020 | B.1.2 | Iowa, USA |
| hCoV-19/deer/USA/IA-208114/2020 | White-Tailed Deer | 12/10/2020 | B.1.2 | Iowa, USA |
| hCoV-19/deer/USA/IA-208115/2020 | White-Tailed Deer | 12/10/2020 | B.1.2 | Iowa, USA |
| hCoV-19/deer/USA/IA-208116/2020 | White-Tailed Deer | 12/10/2020 | B.1.2 | Iowa, USA |
| hCoV-19/deer/USA/IA-208117/2020 | White-Tailed Deer | 12/23/2020 | B.1.2 | Iowa, USA |
| hCoV-19/deer/USA/IA-208329/2020 | White-Tailed Deer | 11/17/2020 | B.1.2 | Iowa, USA |
| hCoV-19/deer/USA/IA-208330/2020 | White-Tailed Deer | 11/17/2020 | B.1.2 | Iowa, USA |
| hCoV-19/deer/USA/IA-208331/2020 | White-Tailed Deer | 11/18/2020 | B.1.1 | Iowa, USA |
| hCoV-19/deer/USA/IA-208334/2020 | White-Tailed Deer | 12/17/2020 | B.1.2 | Iowa, USA |
| hCoV-19/deer/USA/IL-21-038000-001/2021 | White-Tailed Deer | 11/20/2021 | AY.75 | Illinois, USA |
| hCoV-19/deer/USA/IL-21-038000-002/2021 | White-Tailed Deer | 11/21/2021 | AY.3 | Illinois, USA |
| hCoV-19/deer/USA/KS-21-038765-001/2021 | White-Tailed Deer | 12/6/2021 | AY.3 | Kansas, USA |
| hCoV-19/deer/USA/KS-21-038765-002/2021 | White-Tailed Deer | 12/3/2021 | AY.3 | Kansas, USA |
| hCoV-19/deer/USA/MA-21-038766-001/2021 | White-Tailed Deer | 12/1/2021 | AY.119 | Massachusetts, USA |
| hCoV-19/deer/USA/MA-21-038766-002/2021 | White-Tailed Deer | 11/30/2021 | AY.119 | Massachusetts, USA |
| hCoV-19/deer/USA/ME-22-000886-001/2021 | White-Tailed Deer | 12/10/2021 | AY.25 | Maine, USA |
| Tree node ID | **Host** | **Collection date** | **Lineage** | **Location** |
| hCoV-19/deer/USA/ME-22-000886-002/2021 | White-Tailed Deer | 12/11/2021 | AY.103 | Maine, USA |
| hCoV-19/deer/USA/MN-22-000885-001/2021 | White-Tailed Deer | 12/7/2021 | AY.103 | Minnesota, USA |
| hCoV-19/deer/USA/MN-22-000885-002/2021 | White-Tailed Deer | 12/1/2021 | AY.44 | Minnesota, USA |
| hCoV-19/deer/USA/NC-21-038002-001/2021 | White-Tailed Deer | 11/20/2021 | AY.103 | North Carolina, USA |
| hCoV-19/deer/USA/NC-21-038002-002/2021 | White-Tailed Deer | 11/20/2021 | AY.100 | North Carolina, USA |
| hCoV-19/deer/USA/NJ-22-000884-001/2021 | White-Tailed Deer | 12/2/2021 | AY.25 | New Jersey, USA |
| hCoV-19/deer/USA/NJ-22-000884-002/2021 | White-Tailed Deer | 12/8/2021 | AY.42 | New Jersey, USA |
| hCoV-19/deer/USA/NY-21-038001-001/2021 | White-Tailed Deer | 11/24/2021 | P.1 | New York, USA |
| hCoV-19/deer/USA/NY-21-038001-002/2021 | White-Tailed Deer | 11/20/2021 | AY.98.1 | New York, USA |
| hCoV-19/deer/USA/OH-OSU-0025/2021 | White-Tailed Deer | 1/26/2021 | B.1.582 | Ohio, USA |
| hCoV-19/deer/USA/OH-OSU-0057/2021 | White-Tailed Deer | 1/28/2021 | B.1.582 | Ohio, USA |
| hCoV-19/deer/USA/OH-OSU-0058/2021 | White-Tailed Deer | 1/28/2021 | B.1.2 | Ohio, USA |
| hCoV-19/deer/USA/OH-OSU-0078/2021 | White-Tailed Deer | 2/1/2021 | B.1.2 | Ohio, USA |
| hCoV-19/deer/USA/OH-OSU-0079/2021 | White-Tailed Deer | 2/1/2021 | B.1.2 | Ohio, USA |
| hCoV-19/deer/USA/OH-OSU-0109/2021 | White-Tailed Deer | 2/2/2021 | B.1.2 | Ohio, USA |
| hCoV-19/deer/USA/OH-OSU-0141/2021 | White-Tailed Deer | 2/16/2021 | B.1.2 | Ohio, USA |
| hCoV-19/deer/USA/OH-OSU-0212/2021 | White-Tailed Deer | 2/24/2021 | B.1.596 | Ohio, USA |
| hCoV-19/deer/USA/OH-OSU-0335/2021 | White-Tailed Deer | 2/25/2021 | B.1.596 | Ohio, USA |
| hCoV-19/deer/USA/OH-OSU-0336/2021 | White-Tailed Deer | 2/25/2021 | B.1.596 | Ohio, USA |
| hCoV-19/deer/USA/OH-OSU-0340/2021 | White-Tailed Deer | 2/25/2021 | B.1.596 | Ohio, USA |
| hCoV-19/deer/USA/OH-OSU-0341/2021 | White-Tailed Deer | 2/25/2021 | B.1.596 | Ohio, USA |
| hCoV-19/deer/USA/OH-OSU-0343/2021 | White-Tailed Deer | 2/25/2021 | B.1.596 | Ohio, USA |
| hCoV-19/deer/USA/OH-OSU-0344/2021 | White-Tailed Deer | 2/25/2021 | B.1.596 | Ohio, USA |
| hCoV-19/deer/USA/OK-21-038767-001/2021 | White-Tailed Deer | 11/20/2021 | AY.25 | Oklahoma, USA |
| hCoV-19/deer/USA/OK-21-038767-002/2021 | White-Tailed Deer | 11/20/2021 | AY.25 | Oklahoma, USA |
| hCoV-19/deer/USA/PA-21-038768-001/2021 | White-Tailed Deer | 11/28/2021 | AY.107 | Pennsylvania, USA |
| hCoV-19/deer/USA/PA-21-038768-002/2021 | White-Tailed Deer | 11/28/2021 | B.1.1.7 | Pennsylvania, USA |
| Tree node ID | **Host** | **Collection date** | **Lineage** | **Location** |
| hCoV-19/deer/USA/TN-22-000882-001/2021 | White-Tailed Deer | 12/4/2021 | AY.100 | Tennessee, USA |
| hCoV-19/deer/USA/TN-22-000882-002/2021 | White-Tailed Deer | 12/4/2021 | AY.47 | Tennessee, USA |
| hCoV-19/deer/USA/VA-22-000883-001/2021 | White-Tailed Deer | 12/11/2021 | AY.100 | Virginia, USA |
| hCoV-19/deer/USA/VA-22-000883-002/2021 | White-Tailed Deer | 12/11/2021 | AY.42 | Virginia, USA |
| hCoV-19/env/Austria/CeMM21603/2021 | Environment - Composite wastewater sample | 12/28/2021 | BA.1.1 | Austria, UK |
| hCoV-19/env/Austria/CeMM21831/2022 | Environment - Composite wastewater sample | 1/2/2022 | BA.1 | Austria, UK |
| hCoV-19/env/Austria/CeMM21844/2022 | Environment - Composite wastewater sample | 1/2/2022 | BA.1 | Austria, UK |
| hCoV-19/env/Austria/CeMM21850/2022 | Environment - Composite wastewater sample | 1/2/2022 | BA.1.1 | Austria, UK |
| hCoV-19/env/Austria/CeMM21877/2022 | Environment - Composite wastewater sample | 1/1/2022 | BA.1.1 | Austria, UK |
| hCoV-19/env/Austria/CeMM22297/2022 | Environment - Composite wastewater sample | 1/9/2022 | BA.1.1 | Austria, UK |
| hCoV-19/hamster/Hong Kong | Hamster |  | AY.47 | Hong Kong |
| hCoV-19/Syrian hamster/India/MH-ICMR-MCL-21-11828/2021 | Hamster | 11/1/2021 | BA.1.1 | India |
| hCoV-19/USA/NY-NYCPHL-008661/2021 | Human | 12/22/2021 | BA.1.1 | New York, USA |
| hCoV-19/USA/NY-NYCPHL-008664/2021 | Human | 12/23/2021 | BA.1.1 | New York, USA |
| hCoV-19/USA/NY-NYCPHL-008691/2021 | Human | 12/26/2021 | BA.1 | New York, USA |
| hCoV-19/USA/NY-NYCPHL-009078/2021 | Human | 12/29/2021 | BA.1 | New York, USA |
| hCoV-19/USA/NY-NYCPHL-009080/2021 | Human | 12/29/2021 | BA.1.1 | New York, USA |
| hCoV-19/USA/NY-NYCPHL-009082/2021 | Human | 12/30/2021 | BA.1 | New York, USA |
| hCoV-19/USA/NY-NYCPHL-009085/2021 | Human | 12/30/2021 | BA.1 | New York, USA |
| hCoV-19/USA/NY-NYCPHL-009086/2021 | Human | 12/30/2021 | BA.1 | New York, USA |
| hCoV-19/USA/NY-NYCPHL-009087/2021 | Human | 12/31/2021 | BA.1.1 | New York, USA |
| hCoV-19/USA/NY-NYCPHL-009088/2021 | Human | 12/31/2021 | BA.1 | New York, USA |
| hCoV-19/USA/NY-NYCPHL-009090/2021 | Human | 12/31/2021 | BA.1.1 | New York, USA |
| Tree node ID | **Host** | **Collection date** | **Lineage** | **Location** |
| hCoV-19/USA/NY-NYCPHL-009091/2022 | Human | 1/1/2022 | BA.1.1 | New York, USA |
| hCoV-19/USA/NY-NYCPHL-009093/2022 | Human | 1/1/2022 | BA.1 | New York, USA |
| hCoV-19/USA/NY-NYCPHL-009094/2022 | Human | 1/1/2022 | BA.1 | New York, USA |
| hCoV-19/USA/NY-NYCPHL-009095/2022 | Human | 1/1/2022 | BA.1 | New York, USA |
| hCoV-19/USA/NY-NYCPHL-009096/2022 | Human | 1/1/2022 | BA.1 | New York, USA |
| hCoV-19/USA/NY-NYCPHL-009097/2022 | Human | 1/1/2022 | BA.1 | New York, USA |
| hCoV-19/USA/NY-NYCPHL-009098/2022 | Human | 1/1/2022 | BA.1 | New York, USA |
| hCoV-19/USA/NY-NYCPHL-009100/2022 | Human | 1/2/2022 | BA.1 | New York, USA |
| hCoV-19/USA/NY-NYCPHL-009126/2022 | Human | 1/2/2022 | BA.1 | New York, USA |
| hCoV-19/USA/NY-NYCPHL-009127/2022 | Human | 1/2/2022 | BA.1 | New York, USA |
| hCoV-19/USA/NY-NYCPHL-009128/2022 | Human | 1/2/2022 | BA.1.1 | New York, USA |
| hCoV-19/USA/NY-NYCPHL-009129/2022 | Human | 1/3/2022 | BA.1.1 | New York, USA |
| hCoV-19/USA/NY-NYCPHL-009303/2022 | Human | 1/3/2022 | BA.1.1 | New York, USA |
| hCoV-19/USA/NY-NYCPHL-009316/2022 | Human | 1/4/2022 | BA.1 | New York, USA |
| hCoV-19/USA/NY-NYCPHL-009317/2022 | Human | 1/4/2022 | BA.1 | New York, USA |
| hCoV-19/USA/NY-NYCPHL-009318/2022 | Human | 1/4/2022 | BA.1 | New York, USA |
| hCoV-19/USA/NY-NYCPHL-009320/2022 | Human | 1/4/2022 | BA.1 | New York, USA |
| hCoV-19/USA/NY-NYCPHL-009321/2022 | Human | 1/4/2022 | BA.1 | New York, USA |
| hCoV-19/USA/NY-NYCPHL-009323/2022 | Human | 1/5/2022 | BA.1 | New York, USA |
| hCoV-19/USA/NY-NYCPHL-009464/2022 | Human | 1/4/2022 | BA.1 | New York, USA |
| hCoV-19/USA/NY-NYCPHL-009466/2022 | Human | 1/5/2022 | BA.1 | New York, USA |
| hCoV-19/USA/NY-NYCPHL-009468/2022 | Human | 1/5/2022 | BA.1 | New York, USA |
| hCoV-19/USA/NY-NYCPHL-009469/2022 | Human | 1/5/2022 | BA.1 | New York, USA |
| hCoV-19/USA/NY-NYCPHL-009470/2022 | Human | 1/6/2022 | BA.1 | New York, USA |
| hCoV-19/USA/NY-NYCPHL-009471/2022 | Human | 1/3/2022 | BA.1.1 | New York, USA |
| hCoV-19/USA/NY-NYCPHL-009522/2022 | Human | 1/6/2022 | BA.1.1 | New York, USA |
| hCoV-19/USA/NY-NYCPHL-009523/2022 | Human | 1/6/2022 | BA.1.1 | New York, USA |
| Tree node ID | **Host** | **Collection date** | **Lineage** | **Location** |
| hCoV-19/USA/NY-NYCPHL-009529/2022 | Human | 1/7/2022 | BA.1 | New York, USA |
| hCoV-19/USA/NY-NYCPHL-009530/2022 | Human | 1/7/2022 | BA.1.1 | New York, USA |
| hCoV-19/USA/NY-NYCPHL-009532/2022 | Human | 1/7/2022 | BA.1 | New York, USA |
| hCoV-19/USA/NY-NYCPHL-009535/2022 | Human | 1/8/2022 | BA.1.1 | New York, USA |
| hCoV-19/USA/NY-NYCPHL-009538/2022 | Human | 1/5/2022 | BA.1 | New York, USA |
| hCoV-19/USA/NY-NYCPHL-009542/2022 | Human | 1/8/2022 | BA.1 | New York, USA |
| hCoV-19/USA/NY-NYCPHL-009543/2022 | Human | 1/8/2022 | BA.1 | New York, USA |
| hCoV-19/USA/NY-NYCPHL-009545/2022 | Human | 1/9/2022 | BA.1.1 | New York, USA |
| hCoV-19/USA/NY-NYCPHL-009659/2022 | Human | 1/9/2022 | BA.1 | New York, USA |
| hCoV-19/USA/NY-NYCPHL-009660/2022 | Human | 1/10/2022 | BA.1 | New York, USA |
| hCoV-19/USA/NY-NYCPHL-009661/2022 | Human | 1/10/2022 | BA.1.1 | New York, USA |
| hCoV-19/USA/NY-NYCPHL-009662/2022 | Human | 1/10/2022 | BA.1 | New York, USA |
| hCoV-19/USA/NY-NYCPHL-009663/2022 | Human | 1/10/2022 | BA.1.1 | New York, USA |
| hCoV-19/USA/NY-NYCPHL-009664/2022 | Human | 1/10/2022 | BA.1 | New York, USA |
| hCoV-19/USA/NY-NYCPHL-009665/2022 | Human | 1/10/2022 | BA.1.1 | New York, USA |
| hCoV-19/USA/NY-NYCPHL-009666/2022 | Human | 1/7/2022 | BA.1 | New York, USA |
| hCoV-19/USA/NY-NYCPHL-009668/2022 | Human | 1/8/2022 | BA.1.1 | New York, USA |
| hCoV-19/USA/NY-NYCPHL-009669/2022 | Human | 1/8/2022 | BA.1.1 | New York, USA |
| hCoV-19/USA/NY-NYCPHL-009670/2022 | Human | 1/9/2022 | BA.1 | New York, USA |
| hCoV-19/USA/NY-NYCPHL-009671/2022 | Human | 1/9/2022 | BA.1 | New York, USA |
| hCoV-19/USA/NY-NYCPHL-009673/2022 | Human | 1/9/2022 | BA.1.1 | New York, USA |
| hCoV-19/USA/NY-NYCPHL-009685/2022 | Human | 1/10/2022 | BA.1.1 | New York, USA |
| hCoV-19/USA/NY-NYCPHL-009688/2022 | Human | 1/10/2022 | BA.1 | New York, USA |
| hCoV-19/USA/NY-NYCPHL-009692/2022 | Human | 1/11/2022 | BA.1 | New York, USA |
| hCoV-19/USA/NY-NYCPHL-009694/2022 | Human | 1/11/2022 | BA.1.1 | New York, USA |

*The four newly sequenced white-tailed deer origin SARS-CoV-2 genomes from this study were included, together with a total of 135 white-tailed deer origin SARS-CoV-2 isolates (available in GISAID) as well as 63 arbitrarily selected representative human-origin Omicron SARS-CoV-2 genomes circulating in New York City between December 2021 and January 2022. An additional six environmental (wastewater) origin and two reported hamster origin SARS-CoV-2 isolates were also included.
